# Preparatory encoding of the fine scale of human spatial attention

**DOI:** 10.1101/061259

**Authors:** Bradley Voytek, Jason Samaha, Camarin E. Rolle, Zachery Greenberg, Navdeep Gill, Shai Porat, Tahim Kader, Sabahat Rahman, Rick Malzyner, Adam Gazzaley

## Abstract

Our attentional focus is constantly shifting: in one moment our vision may be intently concentrated on a specific spot, while in another moment we might spread our attention more broadly. While much is known about the mechanisms by which we shift our visual attention from place to place, relatively little is know about how we shift the aperture of attention from more narrowly-to more broadly-focused. Here we introduce a novel attentional distribution task to examine the neural mechanisms underlying this process. In this task, participants are presented with an informative cue that indicates the location of an upcoming target. This cue can be perfectly predictive of the exact target location, or it can indicate—with varying degrees of certainty—approximately where the target might appear. This cue is followed by a preparatory period in which there is nothing on the screen except a central fixation cross. Using scalp EEG, we examined neural activity during this preparatory period. We find that with decreasing certainty regarding the precise location of the impending target, participant response times increased while target identification accuracy decreased. Additionally, N1 amplitude in response to the cue parametrically increased with spatial certainty while the multivariate pattern of preparatory period visual cortical alpha (8-12 Hz) activity encoded attentional distribution. Both of these electrophysiological parameters were predictive of behavioral performance nearly one second later. These results offer insight into the neural mechanisms underlying how we use information to guide our attentional distribution, and how that influences behavior.

**Authors contributions:** B.V. and A.G. conceived of the study; B.V. and A.G. designed the experimental task; B.V. and J.S. analyzed the EEG data; B.V., J.S., Z.G., N.G., S.P., T.K., S.R., and R.M. collected and analyzed behavioral data; all co-authors assisted in writing the manuscript.

B.V. is funded by an NIH IRACDA (Institutional Research and Academic Career Development Award), a University of California Presidential Postdoctoral Fellowship, the University of California, San Diego CalIt2 Strategic Research Opportunities Program, and a Sloan Research Fellowship. A.G. is funded by the National Institutes of Health Grant R01-AG30395.

**Significant Statement:** Animals—including humans—frequently shift their visual attentional focus more narrowly or broadly depending on expectations. For example, a predator feline may focus their visual attention on a burrow hole, waiting for their prey to emerge. In contrast, a grizzly bear hunting salmon doesn't know precisely where the fish will jump out of the water, so it must spread its attention more broadly. In a series of novel experiments, we show that this broadening of attention comes at a behavioral cost. We find that multivariate changes in preparatory visual cortical oscillatory alpha (8-12 Hz) encode attentional distribution. These results shed light on the potential neural mechanisms by which preparatory information is used to guide attentional focus.

## Introduction

Humans and animals alike have the ability to prepare for future events and to focus their attention on the spatial location where they expect to observe the upcoming event of interest. Just as a feline stalking its prey can wait patiently—attention focused on a single spot in a clearing or broadly across an entire glade—so too can humans willfully decide to either pay attention to a precise location or spread their attention across their visual field. However, it is well established that performance is worse when attention is distributed compared to when attention is focused, which has been documented as a decrement in performance when we are not given precise details as to the location of the ensuing event (Mangun and Hillyard, 1988) or for cued/attended compared to noncued/unattended locations (Posner, 1980; Shulman et al., 1985; Handy et al., 1996). Attentional cueing is so effective, it can even reduce the effect of hemispatial neglect symptoms (Riddoch and Humphreys, 1983). However, studies that examine the distribution of spatial attention often do so by splitting attention across multiple distinct points in the visual field, such as for multiple object tracking (Cavanagh and Alvarez, 2005; Shim et al., 2013), or by manipulating the distance between the cued location and the upcoming target (Hollingworth et al., 2012). Although a great deal of research has examined the control of focused spatial attention, the neural mechanisms involved in preparatory attentional distribution are less studied.

Here we examined the neural basis for top-down, preparatory spatial attentional focus and distribution using a novel preparatory distributed attention task. In this task, participants are cued as to exactly how narrowly or broadly they need to focus or spread their attention in space in order to detect an impending visual target (Fig. 1 and Methods). We hypothesized that decreased spatial information would both diminish target detection accuracy and increase response time (RT). Moreover, because attention elicits strong modulation of early visual P1/N1 event-related potentials (ERP) (Mangun and Hillyard, 1988; Luck et al., 2000), we expected that the behavioral changes predicted for attentional distribution may be associated with decreased cue-evoked P1 and/or N1 amplitudes related to the reduction in certainty provided by the cue. Furthermore, we predicted that the spatial extent of top-down preparatory attentional distribution will be encoded by the multivariate pattern of later preparatory visual cortical alpha (8-12 Hz) amplitude, allowing us to estimate the attentional focus. This hypothesis is predicated on the idea that, for spatially focused attention versus distributed attention, relatively fewer neurons have to be modulated in a preparatory fashion. This would mean that as the total spatial area to be attended to increases, so too does the number of visual cortical ensembles being modulated. However, this top-down modulation of more visual cortical ensembles would result in a decrement in the precision of attentional distribution, resulting in behavioral performance costs. It is important to emphasize that the visual cortical activity to be analyzed will be during the preparatory period when there is no visual stimulus actually present on the screen other than the central fixation cross; that is, all of the ERP and alpha activity to be analyzed will be preparatory, rather than target-related, allowing us to assess the fine scale of human spatial attention in a manner not possible through behavioral analysis alone.

**Figure 1.**
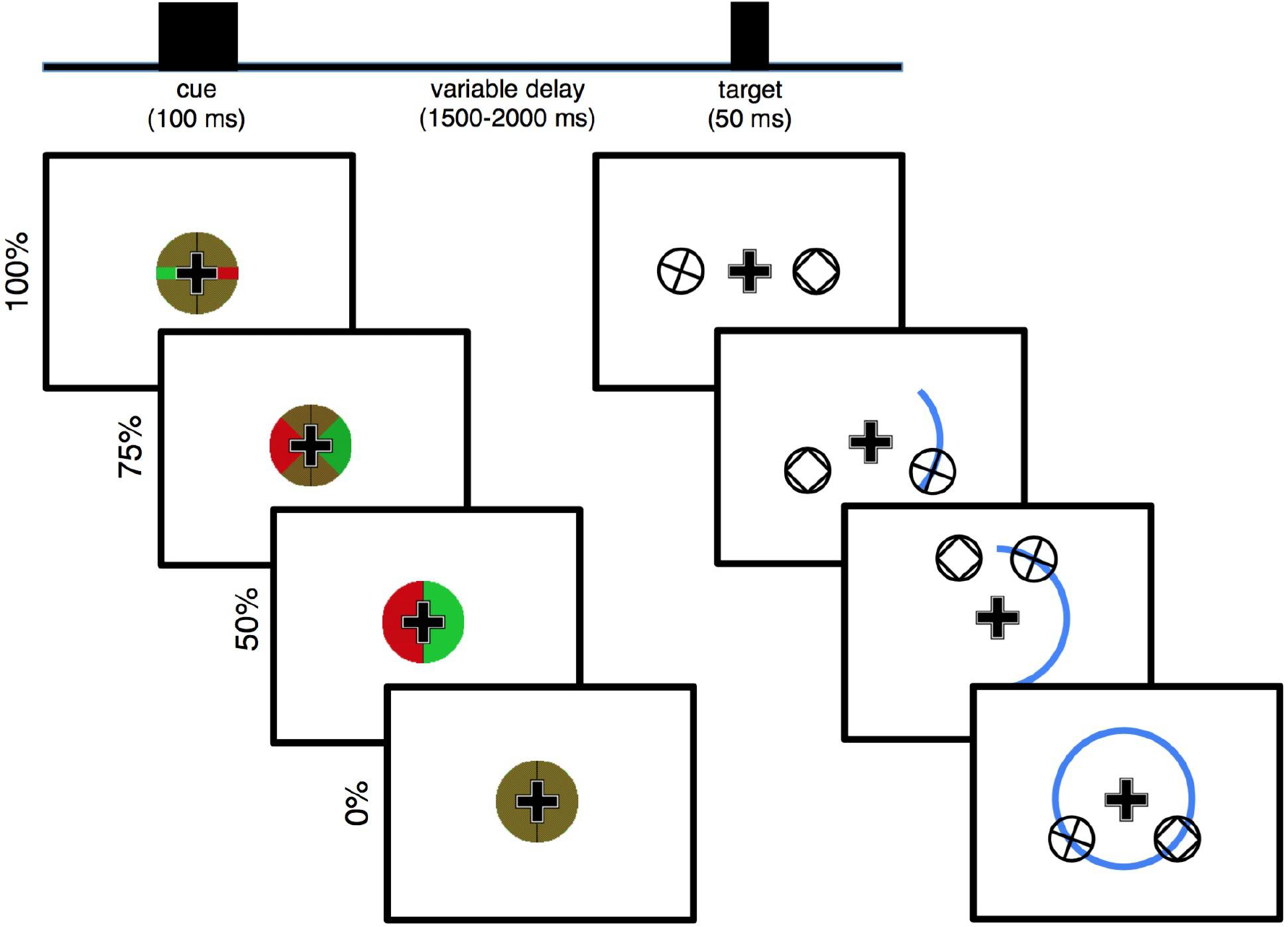
Distributed spatial attention task. Each trial begins with the presentation of a centrally presented spatial cue (left) overlaid on top of a persistent central fixation cross. The cue indicates (with a green wedge) where an upcoming target will appear, with varying certainty, after a random length preparatory period. In the 100% certain condition (top) the target will briefly (50 ms) appear exactly 4.5° to the left or to the right of center (right). In the 75% certainty condition, the target will appear anywhere in a 90° arc, with 4.5° central eccentricity. In the 50% certainty condition, the target will appear anywhere in a 180° arc, while in the 0% certainty condition the target will appear anywhere in the full 4.5° central eccentricity circle. Possible target locations are illustrated with the blue arc (not actually shown on screen). For the bilateral task variant, simultaneous to the presentation of the target a non-target stimulus with matched visual properties is always shown in the non-target hemifield, mirrored across the vertical meridian.

## Materials and Methods

All data were analyzed in MATLAB^®^ (R2014b, Natick, MA) using custom scripts. All participants gave informed consent in accordance with our protocols approved by the UCSF Committee on Human Research in the Human Research Protection Program. Participants in all three experiments were between 20 and 30 years old. There were three total experiments: two behavioral-only experiments with 12 and 9 participants in each, followed by an electroencephalography (EEG) experiment with 17 participants included in the final analyses.

*Experimental Task*. We designed a novel attentional cueing task—a modification of the Posner cueing task (Posner, 1980)—to parametrically manipulate the amount of visual spatial information provided by a pre-target visual cue (Fig. 1). Each trial begins with a centrally presented pre-target cue for 100 ms. This is followed by a variable preparatory period (1500-2000 ms, uniformly distributed) wherein the only stimulus on screen is the central fixation cross. This preparatory period is followed by the visual target, which remains on screen for 50 ms. For the two “bilateral” experiments, simultaneous to the visual target there was also a non-target stimulus (see below). Throughout the entire task, participants were asked to maintain central fixation; a fixation cross is persistent onscreen to assist them. This is to reduce anticipatory saccading toward the hemifield of the upcoming target, maximizing visual extrastriate stimulus representation laterality and minimizing non-neural EEG artifacts (*e.g.*, preparatory saccades).

The cue is a green-and red-checkered circle surrounding the fixation cross, with matched luminances for both colors. This circle is bisected along the vertical meridian with a black line. For the 100% certain condition, the green and red checkerboard is broken by a solid red line in one hemifield and a solid green line in the other hemifield, along the horizontal meridian. These green and red lines are the same vertical width as the arms of the fixation cross, and they extend the entire radius of the cue circle. Whether the green segment appears in the left or right visual hemifield (and thus the red line in the opposite hemifield) is randomized. The hemifield of the green line is perfectly informative of the location of the upcoming target stimulus (100% cue certainty), which will appear 4.5° away from center exactly on the horizontal meridian in whichever hemifield the green line points to. For the 75% certain condition, instead of green and red lines, the cue has green and red 90° wedges, centered along the horizontal meridian. In this condition, the hemifield of the green wedge is still perfectly informative of the hemifield in which the upcoming target will appear, however it also indicates that there is some uncertainty as to where exactly it will appear in that hemifield. Specifically, it indicates that the upcoming target will appear somewhere along a 90° arc, also centered across the horizontal meridian, at 4.5° central eccentricity. For the 50% certain condition, the two hemifields of the cue are either all green or all red, indicating that the upcoming target will appear somewhere along the 180° semicircle (a whole hemifield), at 4.5° central eccentricity. For the 0% certain condition, the cue is just a green and red checkerboard, indicating that the upcoming target will appear somewhere along the 360° circle at 4.5° central eccentricity. Condition (100%, 75%, 50%, or 0 % cue certainty) and target hemifield are randomized on a trial-by-trial basis.

The targets are plusses enclosed by a circle. Participants are tasked to indicate, via manual button press with their dominant hand, whether the plus is exactly vertical and horizontal (index finger) or rotated off-angle (middle finger). For the “bilateral” versions of the experiment, a non-target stimulus is simultaneously presented in the opposite hemifield, mirrored across the vertical meridian. This non-target stimulus is a box enclosed by a circle, meaning its basic visual components (two horizontal and two vertical bars enclosed in a circle) are the same as that of targets, but its context is different. These non-target stimuli are included so that the visual input entering the two visual cortices are largely equivalent during both the cue and target periods, allowing us to isolate cognitive/attention EEG activity from purely visual processes.

Prior to the main experiment, each participant underwent individual psychophysical thresholding to normalize accuracy across participants. The thresholding procedure is a two-down, one-up staircase (converging on ~70% accuracy (Leek, 2001)). In this thresholding task, participants are only presented with the 50% certainty cues, initially being shown either a vertical/horizontal “+”, or a 45°-rotated “X”. With every correct trial, the “X” rotates 1.5° closer toward the vertical/horizontal; with every incorrect response it rotates 3.0° away from the vertical/horizontal. The average angle across the final 10 trials, once behavioral asymptote was reached, was used as the final angle for the main experiment. The average angle across participants was 5.85° (range: 2.20° to 11.25°). Three separate experiments were conducted: in the first—the unilateral variant—12 participants saw a version of the task where only a target stimulus was shown, with no non-target presented in the opposite hemifield. In the second—the full version of the task described above, minus the EEG—was given to 9 participants. In each of these two experiments, each participant performed 200 trials (50 trials per cue information condition).

To examine the effect of cue information on the dependent variables (behavioral and electrophysiological), a linear model was fit on a per-subject basis to get a parameter estimate of the within-subjects effect of cue information on the outcome measure. Under the null hypothesis, the distribution of these parameters estimates (which index the linear change in the dependent variable per cue condition) is not significantly different from zero. This was formally assessed using one-sample, two-tailed *t*-tests, with effect sizes reported as Cohen’s *d*.

*Electroencephalography*. The third experiment included EEG recordings collected from 31 young (20-30 year old) adults (though due to very strict inclusion criteria outlined below, only 17 participants are included in the final analysis). EEG data were collected using a BioSemi ActiveTwo 64 channel DC amplifier with 24-bit resolution, sampled at 1024 Hz. In addition to 64 scalp electrodes both horizontal (HEOG) and vertical (VEOG) electrooculograms were recorded at both external canthi and with a left-inferior eye electrode, respectively. Data were referenced offline to the average potential of two mastoid electrodes and analyzed in MATLAB^®^ (R2014b, Natick, MA) using custom scripts and the EEGLAB toolbox (Delorme and Makeig, 2004).

ERP analyses were performed on bandpass-filtered (0.1-20 Hz) data time-locked to the cue onset using a 100-ms pre-stimulus baseline and 700 ms post-cue time window. Only trials where the participant gave a subsequent correct response were included in EEG analysis. Event onset times were based on timing information provided by a photodiode attached to the stimulus presentation monitor to ensure exact timing relative to stimulus presentation. Eyeblink artifacts were removed using independent component analysis (ICA) (Bell and Sejnowski, 1995). Trials where electrode potentials exceeded ±100 μV and trials with saccades (identified using HEOG channels) were excluded from analysis. Because task stimuli were lateralized, all analyses were performed by hemisphere where contralateral stimuli were defined as left hemisphere channels for right hemifield targets and right hemisphere channels for left hemifield targets (and vice versa). For scalp topography plots, electrode potentials were swapped right to left across the midline to normalize electrode locations as though stimuli were always presented in the right visual hemifield, making left hemisphere channels contralateral to the stimulus and right hemisphere channels ipsilateral to it.

For alpha band (8-12 Hz) analyses, the absolute value of the Hilbert transform of alpha bandpass-filtered continuous (eyeblink corrected) EEG were used to extract alpha band analytic amplitudes. Frequency-band analytic amplitude time series were subjected to normal event-related analyses, removing the same incorrect and artifact-contaminated trials as removed from ERP analyses and normalized against a 100 ms baseline. ERP and analytic amplitude analyses were performed using a visual extrastriate ROI (PO3/4, PO7/8, O1/2).

Each EEG participant performed the full task described above for 400 total trials (100 per cue condition) after pre-EEG psychophysical thresholding. Because the neural questions of interest are predicated on the laterality of top-down preparatory modulation of visual extrastriate regions, we used a very strict EEG artifact rejection procedure wherein any trial with any saccade was dropped from subsequent analysis. This resulted in 14 participants being dropped from subsequent analyses due to too few trials in each condition (25 trial minimum cutoff per condition).

*Inverted Encoding Modeling (IEM)*. Because our hypothesis is multivariate in nature - i.e., the scalp pattern of alpha-band activity representing the attended location will systematically become less selective as cue certainty decreases - we applied a multivariate inverted encoding model (IEM) to quantify topographic patterns of alpha activity representing attentional bias. IEMs model the relationship between neural responses and stimulus or task features using predefined basis functions and have been used to reconstruct basic stimulus features during perception and short-term memory (Sprague and Serences, 2013; Wang et al., 2014; Ester et al., 2015). Recent evidence has shown that IEMs can successfully reconstruct the spatial focus of anticipatory attention from alpha-band topographies (Samaha et al., 2016). Here, our approach was to train a model to distinguish left from right attention during the 100% certain condition, when attention was most spatially focused. We then tested the model on the three other cue certainty conditions (75%, 50%, and 0%), reasoning that the patterns of alpha power should become increasingly dissimilar from the 100% certainty pattern as certainty decreased, reducing the model's ability to discriminate left from right. This approach has the further advantage of reducing a distributed pattern of data into a single metric of attentional bias.

We modeled left versus right spatial attention using a basis set of two binary functions (or “channels”), one representing left spatial attention (*e.g.*, [1 0]) and one representing right (*e.g.*, [0 1]). This approach is analogous to a linear decoding analysis of left versus right attention using differences in classifier evidences to quantify attentional bias (Sprague et al., 2015). As input to the model we used the averaged alpha amplitude from 500-700 ms post-cue from all occipital and parietal electrodes (CPz, CP1/2, 3/4, 5/6, TP7/8, Pz, P1/2, P3/4, P5/6, P7/8, P9/10, POz, PO3/4, PO7/8, Oz, O1/2). In the first step, training data from all but one trial (test data) of the 100% cue certainty condition is used in a general linear model of the form:

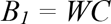

Where *B_1_* (*m* electrodes x *n* trials) is the observed signal at each electrode (alpha amplitude) for each training trial, *C_1_* (*k* channels x *n* trials) is a matrix of predicted responses for each information channel on each trial, and *W* (*m* electrodes x *k* channels) is a weight matrix that characterizes the mapping from “channel space” to “electrode space.” The weight matrix *W* (*m* electrodes x *k* channels) can be derived using ordinary least-squares regression as follows:

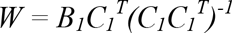

Next, the model is inverted to transform the observed test data *B_2_* (*m* electrodes x 1 trial) into a set of estimated channel responses, *C_2_* (*k* channels x 1 trial), using the weights derived from the training data, via the equation:

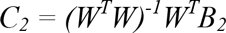

This procedure was iterated until every trial served as a testing set, (*i.e.*, leave-one-trial-out cross-validation). The estimated channel responses were then aligned to a common center and averaged across trials. After each iteration of the cross-validation procedure, the weight matrix *W* was saved. Once cross-validation of the 100% cue certainty condition was completed, these weights were averaged over each iteration and then applied to the independent data from the 75%, 50%, and 0% conditions. Attentional bias (or selectivity) is computed as the subtraction of the output of the channel representing the unattended visual hemifield from that of the attended hemifield. By this metric, zero represents no attentional bias and increasing positive values denote higher channel outputs for the attended hemifield, that is, greater preparatory attentional bias toward the hemifield of the upcoming target.

## Results

The initial behavioral version of the task made use of unilateral stimulus presentation. In this version (*n* = 12 participants), decreasing certainty regarding the spatial location of the upcoming target stimulus reduced participant accuracy and slowed response times (Fig. 2a; acc: *p* = 0.003, *d* = −2.35; RT: *p* =0.011, *d* = 1.84). We then modified the task for EEG to include a non-target stimulus presented simultaneously to the target, but in the opposite visual hemifield (Fig. 1, see Methods). This ensured that visual inputs to both cortical hemispheres were equal across all task conditions as well as during both the cueing and response periods. Behavioral pilot testing of the bilateral design (*n* = 9 participants) revealed the same behavioral pattern: decreasing spatial certainty led to more errors and slower RT (Fig. 2b; acc: *p* = 0.027, *d* = −1.91; RT: *p* = 0.028, *d* = 1.90). Finally, with concomitant EEG recording in another group of participants (*n* = 17) we again observed the same performance pattern, highlighting the robustness of the behavioral effect (Fig. 2c; acc: *p* < 10^−6^, *d* = −4.45; RT: *p* < 10^−7^, *d* = 4.69). Note that only during the EEG recording session were we able to assess eye movements and saccades, with excessive preparatory period saccades resulting in the exclusion of 14 out of 31 total participants from subsequent EEG analyses (leaving *n* = 17). Nevertheless, the 14 excluded participants also showed the same behavioral effect (data not shown; acc: *p* < 10^−5^, *d* = − 4.55; RT: *p* = 0.014, *d* = 1.58).

**Figure 2.**
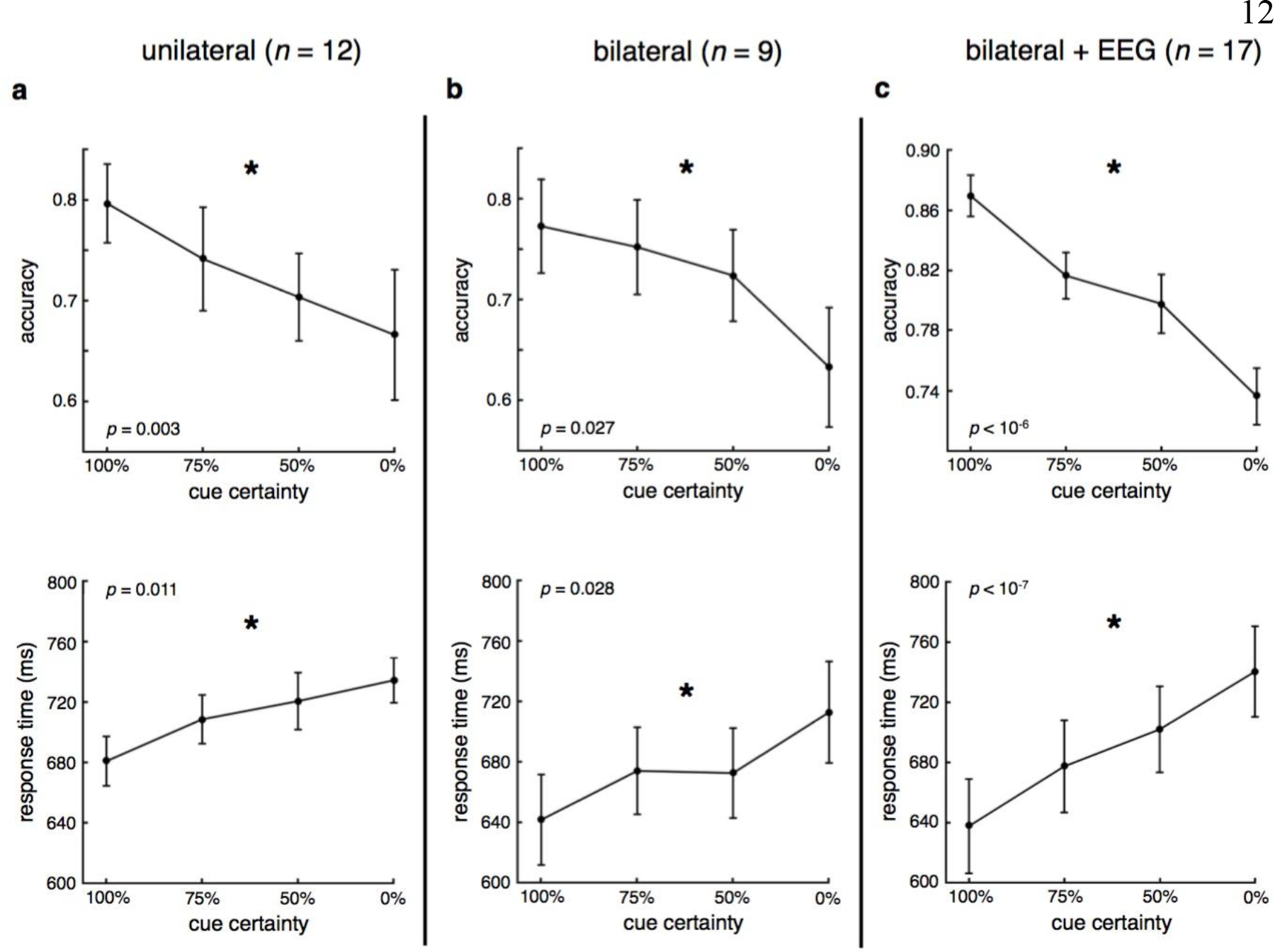
Behavioral results. **a-c**, Across three separate experiments using two variants of the behavioral paradigm, we find that decreasing spatial certainty regarding the location of upcoming target stimulus resulted in less accurate, slower responses. (^*^significant effect of spatial certainty, *p* < 0.05; error bars: sem).

Electrophysiological analysis of cue-evoked visual extrastriate N1 (140-180 ms) activity showed that both contralateral and ipsilateral N1, but not P1, were modulated by cue certainty such that average N1 amplitude magnitude decreased with spatial uncertainty (Fig. 3, top; contralateral: *p* = 0.039, *d* = 1.12; ipsilateral: *p* = 0.058, *d* = 1.02). N1 magnitude also trended toward being greater at contralateral, compared to ipsilateral, sites (*p* = 0.097, *d* = 1.12) and for the high certainty conditions (post-hoc one sample, two-tailed *t*-test for differences from 0 μV, contra-/ipsi-lateral: *p*_*100%*_ = 0.037/0.078, *p*_*75%*_ = 0.028/0.063, *p*_50%_ = 0.50/0.52, *p*_*0%*_ = 0.55/0.79). Note the N1 effect was largest for the 75% condition, with no P1 effect. Given how small the cue was, and how luminance was held constant for the different cue condition stimuli, these relatively weak effects are not surprising.

Analysis of focal, event-related visual extrastriate alpha amplitude showed a strong, early (250-450 ms) alpha amplitude decrease, followed by a sustained alpha negativity (500-700 ms) (Fig. 3, bottom). However, neither early nor late visual extrastriate alpha amplitudes were parametrically modulated by the cue, in either hemifield (early contra: *p* = 0.14, *d* = 0.79; early ipsi: *p* = 0.40, *d* = 0.43; late contra: *p* = 0.70, *d* = 0.20; late ipsi: *p* = 0.71, *d* = −0.19). Note that post hoc analysis of univariate alpha shows that 0% certainty is significantly different from the other three conditions (*p*_*100*_ = 0.024, *p*_*75*_ = 0.033, *p*_*50*_ = 0.088), but it is insensitive to the finer-grained allocation of attention, which may be better captured via the multivariate topography of alpha.

**Figure 3.**
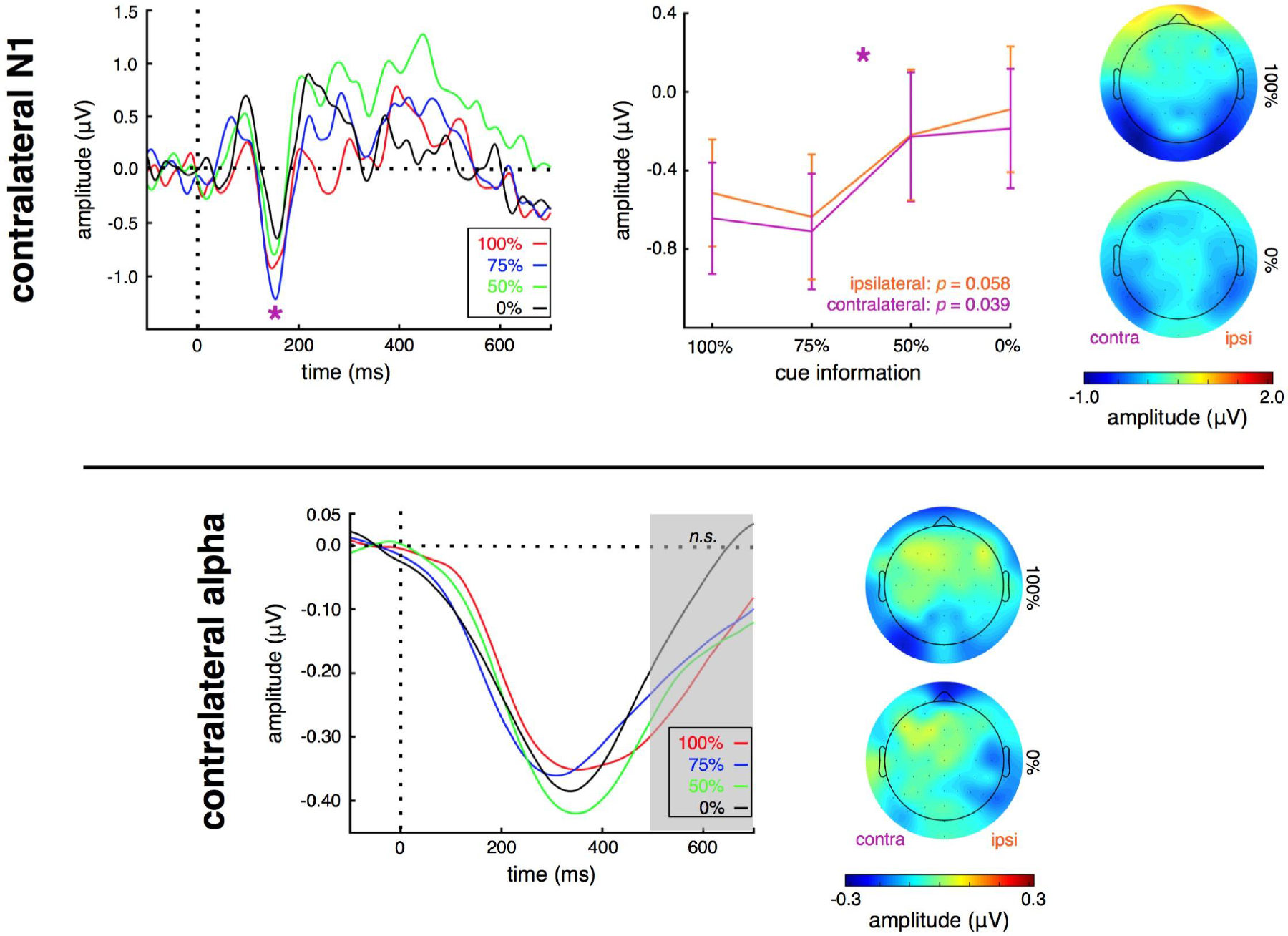
Univariate cue-evoked N1 and alpha responses. (Top) cue-evoked visual extrastriate N1 amplitude is largest for conditions where attention needs to be deployed more focally. (Bottom) In contrast, neither early nor late univariate alpha amplitude is significantly modulated by attention distribution requirements. (^*^significant effect of spatial certainty, *p* < 0.05; *n.s.*: not significant; error bars: sem).

To examine preparatory period visual cortical encoding of attentional distribution, we used an inverted encoding model (IEM) (Sprague et al., 2015) (see Methods). Here, the IEM takes into account the multivariate spatial pattern of late alpha activity across all parietal and occipital sites (see Methods) to get a trial-by-trial estimate of each participant’s attentional bias for each cueing condition (toward or away from the cued location). We find that with decreasing certainty of the upcoming target location, participants showed declining attentional bias (Fig. 4a; *p* = 0.009, *d* = −1.48). This was driven by a significant bias toward the cued location for the 100% condition, with increasingly weaker bias with decreasing certainty (*post-hoc* one sample *t*-test: *p*_*100*_ = 0.038, *p*_*75*_ = 0.10, *p*_*50*_ = 0.10, *p*_*0*_ = 0.35).

A complementary approach to examine the role of the spatial patterning of late visual alpha activity in attentional distribution is to assess trial-by-trial interhemispheric correlations. That is, for each participant, for each condition type, for each trial, we can look at how similar the alpha amplitudes are in the contralateral and ipsilateral hemispheres. Here, the hypothesis is that for more focused conditions there will be greater preparatory, unilateral, top-down modulation of contralateral alpha, leading to relatively weak interhemispheric correlations caused by stronger unilateral modulation. In contrast, for more broadly distributed attention conditions this top-down modulation will be more balanced across both visual hemispheres, leading to stronger interhemispheric correlations. Confirming our hypothesis, we find that as participants prepare to distribute their attention more broadly, trial-by-trial correlation of contralateral and ipsilateral late visual alpha increases (Fig. 4b; *p* = 0.015, *d* = 1.37). This pattern was not observed for N1 amplitude to the cue (*p* = 0.50, *d* = 0.35).

**Figure 4.**
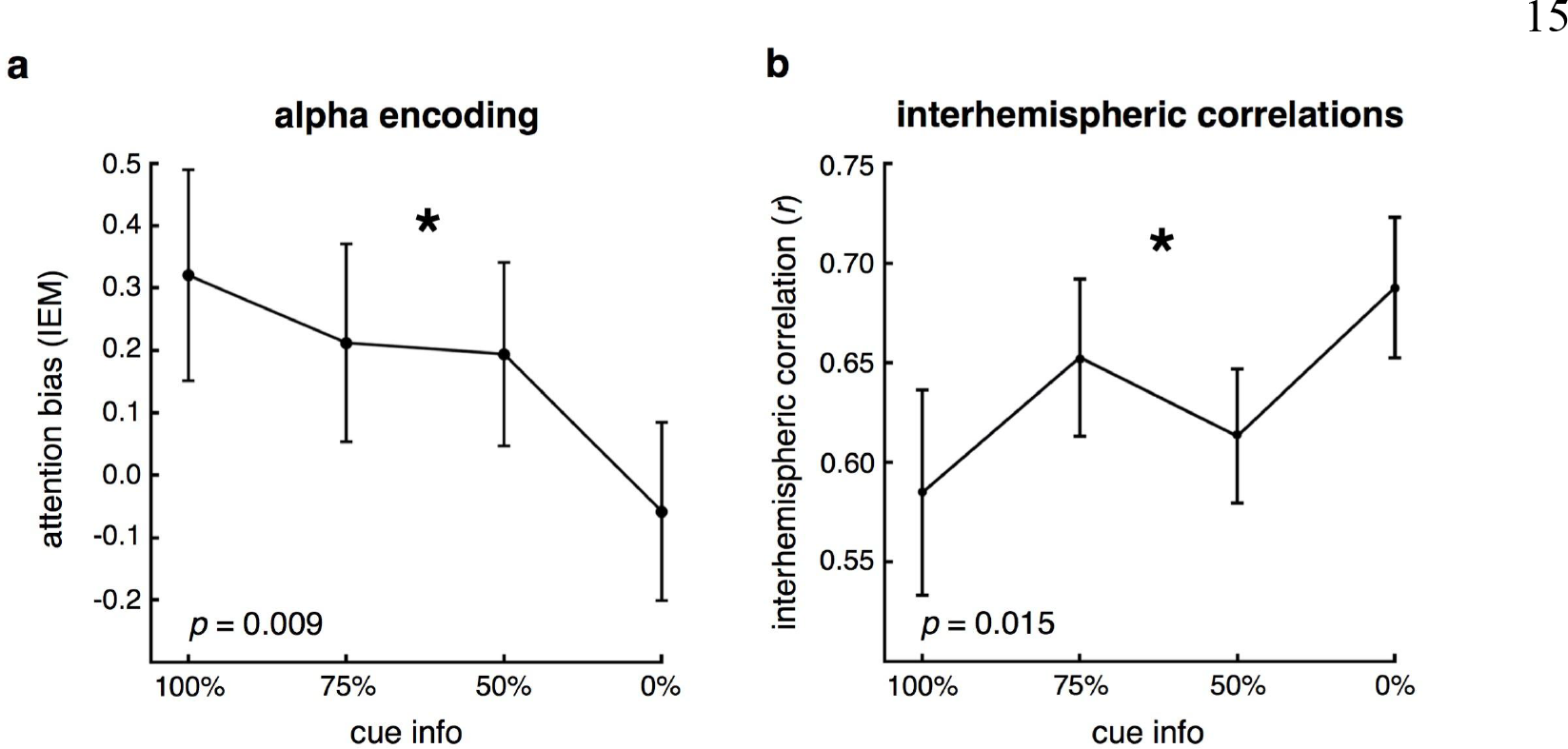
Multivariate alpha encoding and interhemispheric alpha correlations. **a**, Inverted encoding model shows that multivariate alpha spatial patterning in visual extrastriate encodes the bias of attention. Multivariate alpha patterns representing the attended location become less hemifield-selective as spatial attention becomes less focused. This pattern is manifest as a linear decline in IEM bias as a function of cue information, reaching zero bias when the cue is completely uninformative. **b**, In addition, interhemispheric visual alpha amplitudes are relatively less correlated across trials for the focused, 100% conditions, and become increasingly more correlated as attention needs to be distributed more broadly and, ultimately, across hemifields. (^*^significant effect of spatial certainty, *p* < 0.05; error bars: sem).

Finally, we observe that preparatory period electrophysiological activity predicts subsequent behavioral performance. We modeled difference in accuracy or RT between the 100% and 0% certainty conditions as a function of the concomitant difference in contralateral visual N1 amplitude or alpha bias (from the IEM). We find that differences in both N1 amplitude and alpha biasing predicts both accuracy and RT difference (Fig. 5; *r*_*N1/acc*_ = −0.57, *p* = 0.017; *r*_*N1/RT*_ = 0.53, *p* = 0.028; *r*_*aipha/acc*_ = −0.48, *p* = 0.049; *r*_*aipha/RT*_ = 0.50, *p* = 0.021). That is, the participants with the largest N1 amplitude and alpha bias differences between 100% predictive information and 0% information, showed the biggest behavioral differences, characterized by both a greater decrement in accuracy and a slowing of RT.

While N1 amplitude and alpha selectivity differences each predict behavior, independently explaining 28% and 18% of the variance in accuracy difference, respectively, multiple linear regression modeling using both electrophysiological variables as predictors improves prediction of behavioral accuracy to 34% (adjusted *R*^*2*^ to control for the number of predictors in the model, *p* = 0.036). Similarly, N1 and alpha selectivity differences independently explain 24% and 26% of the variance in RT difference, respectively; this improves to 37% when both electrophysiological variables are included (*p* = 0.003; note that N1 difference is not correlated with alpha bias difference: *r* = 0.32, *p* = 0.21). These results suggest that these two electrophysiological variables have relatively independent influences on upcoming behavioral outcomes, and are robust to using slope change as opposed to the 100% versus 0% difference (acc model: adjusted *R*^*2*^ = 0.46, *p* = 0.020; RT model: adjusted *R*^*2*^ = 0.42, *p* = 0.030).

**Figure 5.**
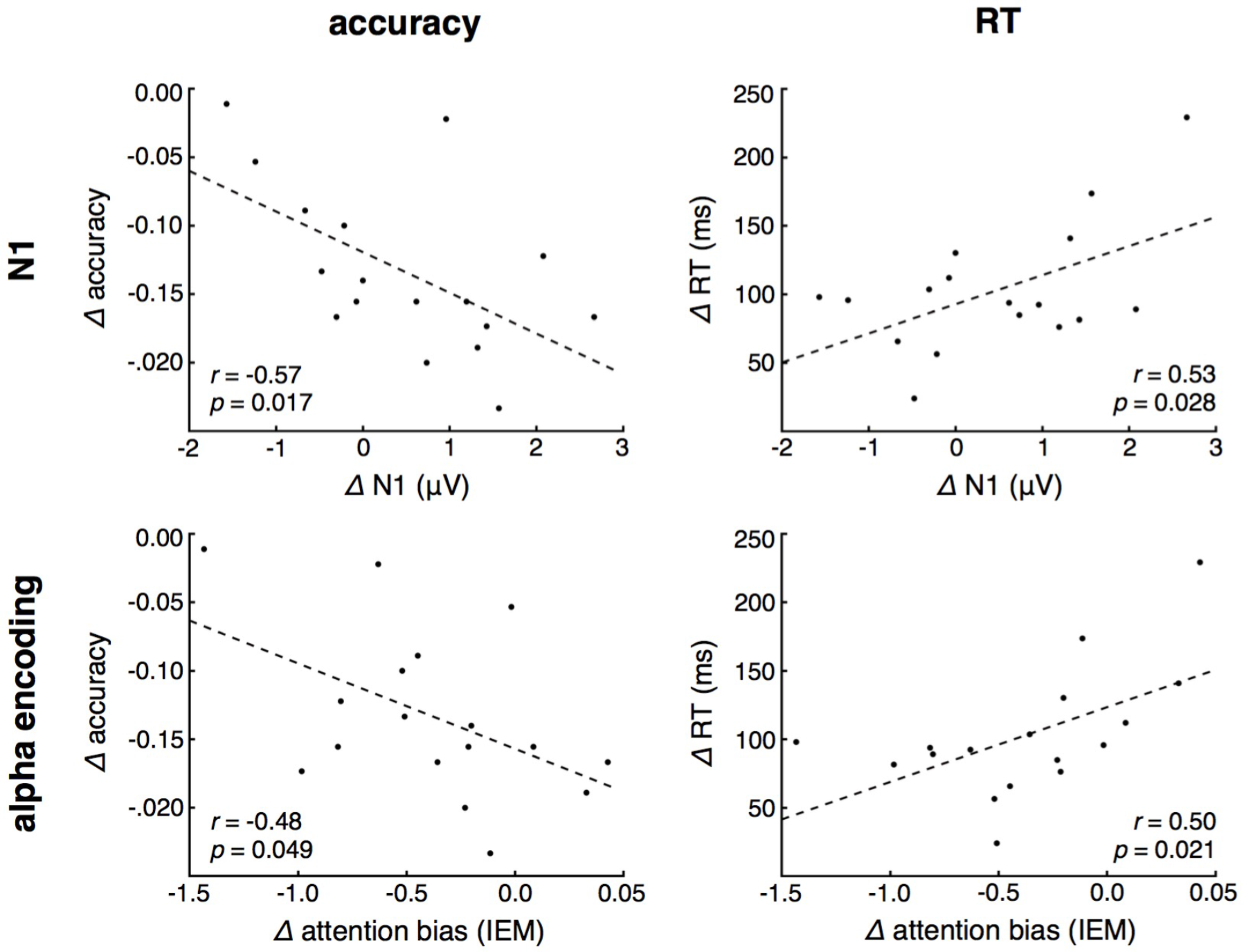
Electrophysiological behavioral prediction. Difference in N1 and alpha encoding from the 100% to 0% conditions predict difference in accuracy and response times. (^*^significant effect of spatial certainty, *p* < 0.05; dashed line: linear fit).

## Discussion

Spatial attention is a critical aspect of cognition, allowing animals to navigate through our complex world and make rapid decisions efficiently and effectively. While a great deal is known regarding how spatial attention is deployed to specific regions of the visual field, how it is used to track and follow objects, and how attentional signals gets passed between the two cerebral hemispheres (Drew and Vogel, 2008), relatively little is understood about how preparatory information is used to focus or spread attention as needed. Previous work has shown that attention to lateralized visual targets modulates early measures of cortical activity, namely the P1 and N1 ERPs (Mangun and Hillyard, 1988; Luck et al., 2000), with unilateral prefrontal cortex (PFC) lesions disrupting these ERPs only for stimuli presented contralateral to the PFC lesion, suggesting that hemispheric control of attention is semi-independent (Barcelo et al., 2000; Battelli et al., 2009; Voytek and Knight, 2010).

Although ERPs provide a robust index of top-down attentional modulation of neural activity in visual extrastriate cortex, the P1 and N1 are short lasting, time-locked signals, and are therefore perhaps less appropriate for assessing preparatory attentional distribution. In contrast, event-related alpha amplitude can be used to assess the degree of lateralized attention (Worden et al., 2000) and is sustained throughout preparatory and delay periods (Palva and Palva, 2007; Capotosto et al., 2009; Jensen and Mazaheri, 2010; Rohenkohl and Nobre, 2011). Physiologically, alpha amplitude is inversely correlated with cortical potentiation (Jasper and Penfield, 1949), making it an ideal index of top-down preparatory modulation of visual cortex (Palva and Palva, 2007).

In order to assess the neural mechanisms underlying the preparatory distribution of attention, we used a novel distributed attention task, combined with scalp EEG, to examine how preparatory period visual cortical alpha activity influences behavioral outcomes more than a second later. The behavioral results suggest that participants are challenged by the task of distributing their attention to broader visual areas, and that when they have to focus their attention to only one location they can respond more quickly and accurately. Participants were only given between 1.5 and 2.0 seconds during the preparatory period to make use of the cue in preparation for the upcoming target.

However, from behavior alone it is unclear whether the accuracy and RT costs associated with more distributed attention are driven purely by a spatial search cost *after* the target appears, or whether participants make use of the cue information during the preparatory period to improve their performance. By focusing EEG analyses on the preparatory period only, when no task-related visual information was present on screen, we were able to isolate a neural mechanism of preparatory attentional distribution.

The fact that cue-evoked contralateral N1 amplitude and multivariate alpha encoding during the preparatory period are both predictive of later behavioral outcomes shows how participants effectively use contextual cues to optimize attentional focus in preparation of a future event. The N1 and alpha reflect different physiological processes, with the former reflecting early visual processing of the initial information-giving cue, and the latter likely indexing top-down visual cortex activity modulation in preparation of the ensuing target. We found that both N1 and alpha independently predict later performance, and that adding both electrophysiological variables into the behavioral prediction models improves the predictions. That is, the behavioral modeling suggests N1 and alpha provide independent information, and thus are likely reflecting different physiological processes needed for successful task performance.

Interestingly, while univariate, focal alpha is different for 0% compared to the more informative cueing conditions, it is insensitive to finer differences between conditions. However, we hypothesized that preparatory attention would not affect just local alpha amplitude, but rather multivariate alpha topography and *amplitude*, captured via the inverted encoding model. The combined N1 and alpha IEM results strongly suggest that the distribution of attention is not solely an attentional search problem where one must find a target within a visual field. Rather, informative cues influence both early visual processing of the cue that supplies the predictive information itself, reflected in the N1, in conjunction with later preparatory spatial attentional deployment, indexed via multivariate alpha distribution. It is important to note that such decoding methods, especially at the level of scalp EEG, are unlikely to capture topographic maps of feature selectivity, for example; rather they are more likely to reflect more coarse-scale maps (Freeman et al., 2011; Wang et al., 2014).

We found that participants who showed the biggest difference in multivariate alpha between the focused (100%) versus distributed (0%) conditions also showed the smallest behavioral difference. In interpreting this result, a small performance difference could either be considered positive if participants performed consistently well, or negative if they were consistently poor performers. Upon further examination, participants with the smallest accuracy difference between 100% and 0% exhibited the highest performance for the uncertain, 0%, condition (*r* = 0.69, *p* = 0.0021). This observation suggests that *failure*— to modulate the multivariate pattern of preparatory visual alpha is associated with poorer overall performance. That is, the behavioral cost associated with distributing attention across broader spatial fields is driven by the inability to modulate the pattern of preparatory visual cortical alpha. Thus, the ability to more precisely modulate the visual cortical neurons, perhaps through gain control mechanisms (Hillyard et al., 1998), that represent visual fields of varying extents improves performance overall across all conditions, reducing the magnitude of performance declines associated with distributed attentional focus. The results show that preparatory attention can be finely tuned and spatially modified rapidly depending on context, which in turn biases the cortex for target detection and influences behavioral outcomes more than a second later.

